# Comparing the XGBoost machine learning algorithm to polygenic scoring for the prediction of intelligence based on genotype data

**DOI:** 10.1101/2022.06.12.495467

**Authors:** Laura Fahey, Derek W. Morris, Pilib Ó Broin

## Abstract

A polygenic score (PGS) is a linear combination of effects from a GWAS that represents and can be used to predict genetic predisposition to a particular phenotype. A key limitation of the PGS method is that it assumes additive and independent SNP effects, when it is known that epistasis (gene interactions) can contribute to complex traits. Machine learning methods can potentially overcome this limitation by virtue of their ability to capture nonlinear interactions in high dimensional data. Intelligence is a complex trait for which PGS prediction currently explains up to 5.2% of the variance, a relatively small proportion of the heritability estimate of 50% obtained from twin studies. Here, we use gradient boosting, a machine learning technique based on an ensemble of weak prediction models, to predict intelligence from genotype data. We found that while gradient boosting did not outperform the PGS method in predicting intelligence based on SNP data, it was capable of achieving similar predictive performance with less than a quarter of the SNPs with the top SNPs identified as being important for predictive performance being biologically meaningful. These results indicate that ML methods may be useful in interpreting the biological meaning underpinning SNP-phenotype associations due to the smaller number of SNPs required in the ML model as opposed to the standard PGS method based on GWAS.

## 1 Introduction

Common complex disorders and traits are mainly influenced by common genetic variants with small effect sizes. The polygenic model of inheritance states that it is the cumulative contribution of many such variants that influences the phenotype (Visscher and Wray, 2015). Genome-wide association studies (GWAS) investigate the contribution of individual variants, most often single nucleotide polymorphisms (SNPs), to the phenotype in question and for each SNP calculates an effect size and a p-value. A polygenic score (PGS) summarizes the genetic effects from GWAS into a value that can be used to predict genetic predisposition to a particular phenotype. There are multiple methods available that compute PGSs from GWAS results. P-value based clumping and thresholding (PC + T) is the most basic method and has been the most frequently used to date (Ni *et al*., 2021). Here, linkage disequilibrium (LD) clumping is performed to obtain an independent set of SNPs, followed by the application of a p-value threshold to select the most informative SNPs (multiple p-values are tested to identify the value that performs the best). A PGS is then calculated for each sample by summing up, for each SNP, the number of minor alleles at that locus multiplied by the GWAS effect size for that SNP. More sophisticated methods, such as LDpred2 (Privé *et al*., 2020) and SBAYESR (Lloyd-Jones *et al*., 2019), that estimate SNP effects in a way that accounts for different genetic architectures using external LD reference panels, have been shown to outperform PC + T (Ni *et al*., 2021), and are therefore becoming more popular.

PGSs have not yet been very successful because they explain only a very small proportion of the predicted heritability from family studies for most complex diseases and traits (Torkamani *et al*., 2018). There are multiple potential reasons for this, including the fact that current methods do not account for different SNPs conferring risk under different genetic models (additive, dominant or recessive) and they also do not account for interactions between SNPs, often referred to as epistasis. Epistasis is difficult to detect on a genome-wide scale. Testing all possible interactions would not be computationally feasible and studies that restricted tests to two loci interactions failed to identify statistically significant associations due to the necessity for conservative significance levels to correct for so many tests (Lippert *et al*., 2013). Machine learning (ML) algorithms that make minimal assumptions about causal mechanisms and can model nonlinear interactions can potentially overcome these limitations, providing a powerful alternative to PGS approaches.

A handful of studies have compared the performance of standard linear PGS analysis to a variety of ML methods including a support vector machine (Behravan *et al*., 2018; Romagnoni *et al*., 2019),gradient boost machines (GBMs) (Behravan *et al*., 2018; Romagnoni *et al*., 2019; Gola *et al*., 2020), a dense neural network (Behravan *et al*., 2018; Romagnoni *et al*., 2019; Gola *et al*., 2020), a random forest (Behravan *et al*., 2018; Romagnoni *et al*., 2019; Gola *et al*., 2020), and an artificial neural network (Pinto *et al*., 2019). These studies looked at a variety of different phenotypes, including breast cancer, Crohn’s disease, coronary artery disease and schizophrenia, and achieved varying results in terms of whether the PGS or ML method performed better; this may be attributable to the fact that genetic interactions between SNPs contribute to some phenotypes but not others (Polderman *et al*., 2015). Alternatively, differences in study design, including different feature selection methods, different ML models, differences in sample size and failure to account for confounders, may also play a role.

Here, we build on recent research by using the GBM algorithm, XGBoost, to predict intelligence, a complex trait for which PGS prediction currently explains 5.2% of the variance (Savage *et al*., 2018), a considerably lower value than the heritability estimate of 50% obtained from twin studies (Polderman *et al*., 2015).

We chose a GBM algorithm due to its ability to identify features of interest, ease of interpretation, speed and consistently impressive performance in Kaggle (https://www.kaggle.com/) machine learning competitions. Furthermore, the process of testing different SNP values at each binary split in each tree makes such tree based algorithms more flexible to different genetic models i.e additive, recessive or dominant (Goldstein *et al*., 2010).

Here, we compare the performance of XGBoost to the standard PGS method in predicting intelligence based on SNP data. We are using the PC + T method for PGS construction as it is more directly comparable to XGBoost for two reasons: 1) It does not require external data, and 2) it uses a subset of SNPs (it is not possible to model all SNPs in XGBoost due to the memory burden). We then compare and contrast the SNPs identified as important by both of these methods as well as the genes the SNPs map to and the biological pathways that they are enriched in.

## 2 Methods

### 2.1 Phenotype and UK Biobank Data Quality Control

In this study we use the Fluid Intelligence score from UK Biobank (UKB) (data-field ID 20016) as a measure of intelligence. The Fluid Intelligence test was completed on site at UKB assessment centers on a touch screen computer. It consists of 13 multiple choice questions assessing verbal (e.g., “If Truda’s mother’s brother is Tim’s sister’s father, what relation is Truda to Tim?” Possible answers: Aunt/Sister/Niece/Cousin) and numerical (e.g. “100…99…95…86…70… What comes next?” Possible answers: 50/49/48/47/46/45) reasoning abilities. The score is the number of correctly answered questions in 2 minutes (consists of 14 discrete values (0-13)) (UK Biobank, 2012). In assessing the validity of UKB cognitive tests, Fawns-Ritchie & Deary, 2020 found that UKB Fluid Intelligence test scores positively correlated with general cognitive ability that was assessed using 11 reference tests (R^2^ = 0.553, *p* < 0.001). Deary *et al*., 2022 pointed out however that the name of the test “Fluid Intelligence” is incorrect and that it should instead be called “Verbal and Numerical Reasoning”, therefore the latter is the name we use in this study.

Genotypic data was collected, processed and imputed by UK Biobank (UKB; Bycroft *et al*., 2018). We restricted samples to those that were unrelated and of European ancestry. Further samples were removed on the basis of high SNP missingness, unusually low or high heterozygosity, discordent sex informaiton, and chromosomal aneuploidies. Imputed variants were converted to hard calls at a certainty threshold of 0.9. SNPs were excluded if their proportion of missing genotypes exceeded 2%, minor allele frequency (MAF) was less than 1%, or Hardy–Weinberg equilibrium (HWE) p-value was lower than 1×10^-6^. Multi-allelic snps and SNPs with differences in allele frequency between the two genotyping arrays used were also removed. The total number of samples and SNPs after QC was 118,021 and 7,832,917, respectively. Further detail on QC methods are provided in the Supplementary Material. Code used is freely available at https://github.com/laurafahey02/UK_Biobank_QC.

We created a binary phenotype by assigning individuals in the first and fifth quantiles, respectively, to the groups “HIGH” and “LOW” (Figure 1). Samples in the HIGH group were considered case (N samples = 31,824) and samples in the LOW group were considered control (N samples = 25,734). This method of phenotype generation reduces the total number of samples available for analyses, but enables us to define more distinct sample groups increasing the likelihood of detecting SNPs contributing to phenotypic differences. To confirm that this phenotype captures the same genetic effect as the full quantitative phenotype, we compared the resulting odds ratios from logistic regression to the published beta values from a GWAS linear regression performed by GWAS ATLAS (https://atlas.ctglab.nl/) (Watanabe *et al*., 2019). As expected, there is a linear trend, with a Pearson correlation of 0.7 (Supplementary Figure 1).

**Figure 1:**
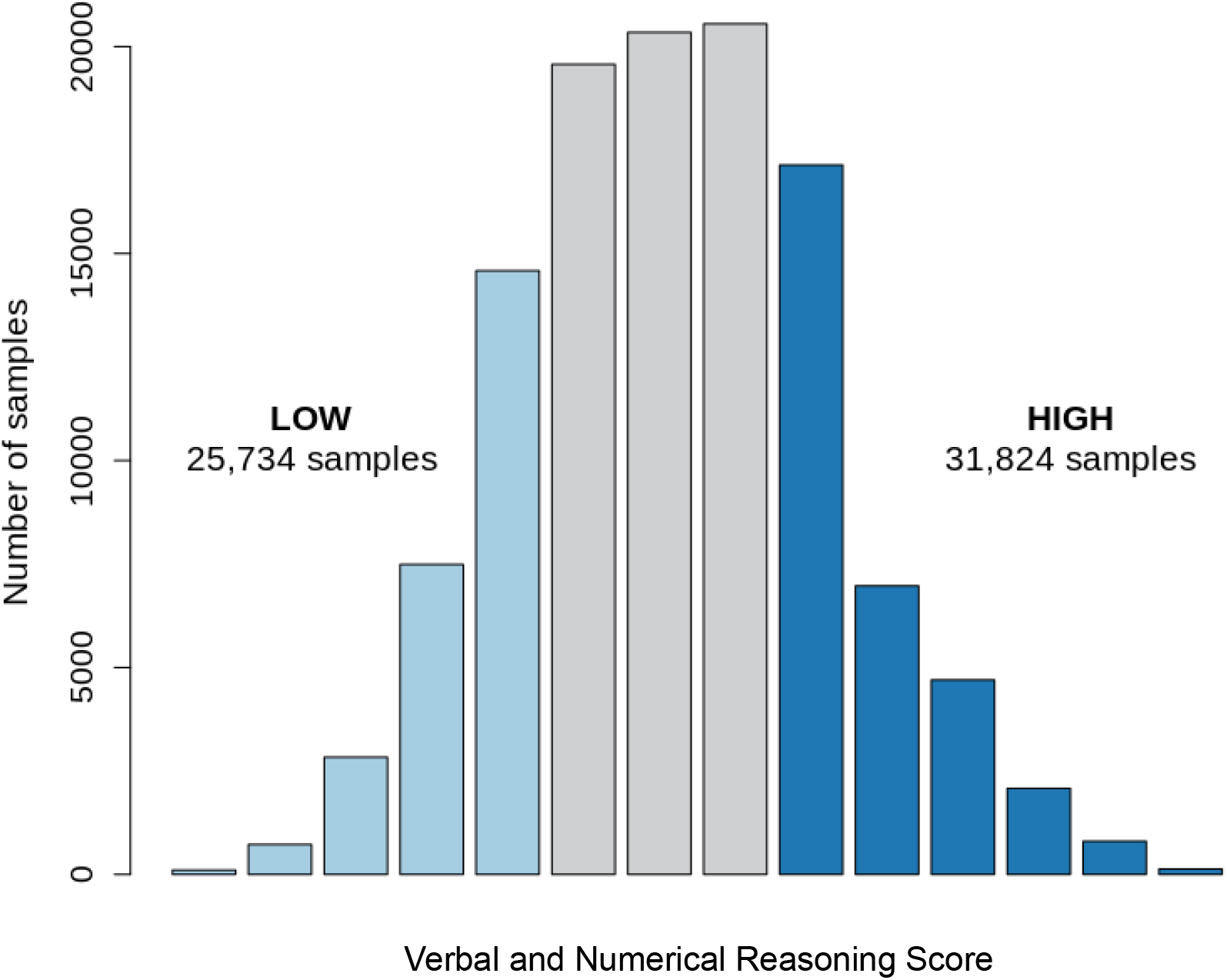
Distribution of Verbal and Numerical Reasoning Scores and assignment to LOW and HIGH groups. Bar chart depicting the number of samples obtaining each verbal and numerical reasoning score (0-13). Bars coloured light blue represent the verbal and numerical reasoning scores of samples in the LOW group and bars coloured dark blue represent the verbal and numerical reasoning scores of samples in the HIGH group. Samples from the gray-coloured bars were not included in this study.

### 2.2 Feature Selection

Feature selection is performed prior to running the XGBoost model in order to avoid the curse of dimensionality (Ross *et al*., 2009) as well as to reduce the computational burden. Here, we used a combination of filtering by SNP annotation, by p-value from a logistic regression GWAS and LD clumping to obtain a set independent, biologically relevant SNPs.

A previously published meta-GWAS of intelligence found enrichment of significant SNPs in conserved, coding and H3K9 histone acetylated regions (markers of active chromatin) (Savage *et al*., 2018). We annotated SNPs to these functional categories using data from the LDSC tool (https://github.com/bulik/ldsc), and restricted our analyses to only these SNPs. This reduced the total number of SNPs in our analysis to 1,099,085.

We then performed a logistic regression GWAS in PLINK2 with firth-fallback and obtained a p-value of association for each SNP. We restricted SNPs to those that were LD-independent, keeping the most significant SNP in each LD block, using clumping (PLINK2 parameters: --clump-;kb 250 --clump-r2 0.1). We next applied a further p-value threshold that was optimized as a hyperparameter of the ML approach.

### 2.3 Encoding

Genotype information was converted from PLINK bed file format to text file format according to an additive genetic model. This was done using the --export A flag in PLINK2 which encodes each genotype as a single dosage number representing the number of minor alleles at that position (values = 0, 1 and 2).

### 2.4 XGBoost Implementation and Workflow

XGBoost was implemented using the scikit-learn machine learning library in Python (https://scikit-learn.org/stable/). Samples were split 80:20, using four folds as training data and one fold as the held out test data to report the final performance. This was done in Python using *Stratifiedkfold* to preserve the original case-control ratio in the train and test subsets. The training data was further split into three folds for three-fold cross-validation (CV) to evaluate the best performing hyperparameters (Figure 2). Hyperparameter optimisation was done using a semi-manual stepwise method. This method was chosen because of it being efficient in terms of time and memory. The first hyperparameter optimized was the GWAS p-value threshold for feature selection. A GWAS was run for each inner training CV fold (three times) and LD clumping was performed. Next, a p-value threshold was applied and varied whilst setting the other hyperparameters to values which had been empirically shown to perform well during initial experiments (hyperparameters:‘colsample_bytree’:0.5,‘colsample_bylevel’: 0.5, ‘colsample_bynode’: 0.5, ‘learning_rate’: 0.05, ‘max_depth’: 20, ‘min_child_weight’: 50, ‘n_estimators’: 15000, ‘subsample’: 0.5). The p-value threshold that resulted in the best performance on the validation subset was used in subsequent steps.

**Figure 2:**
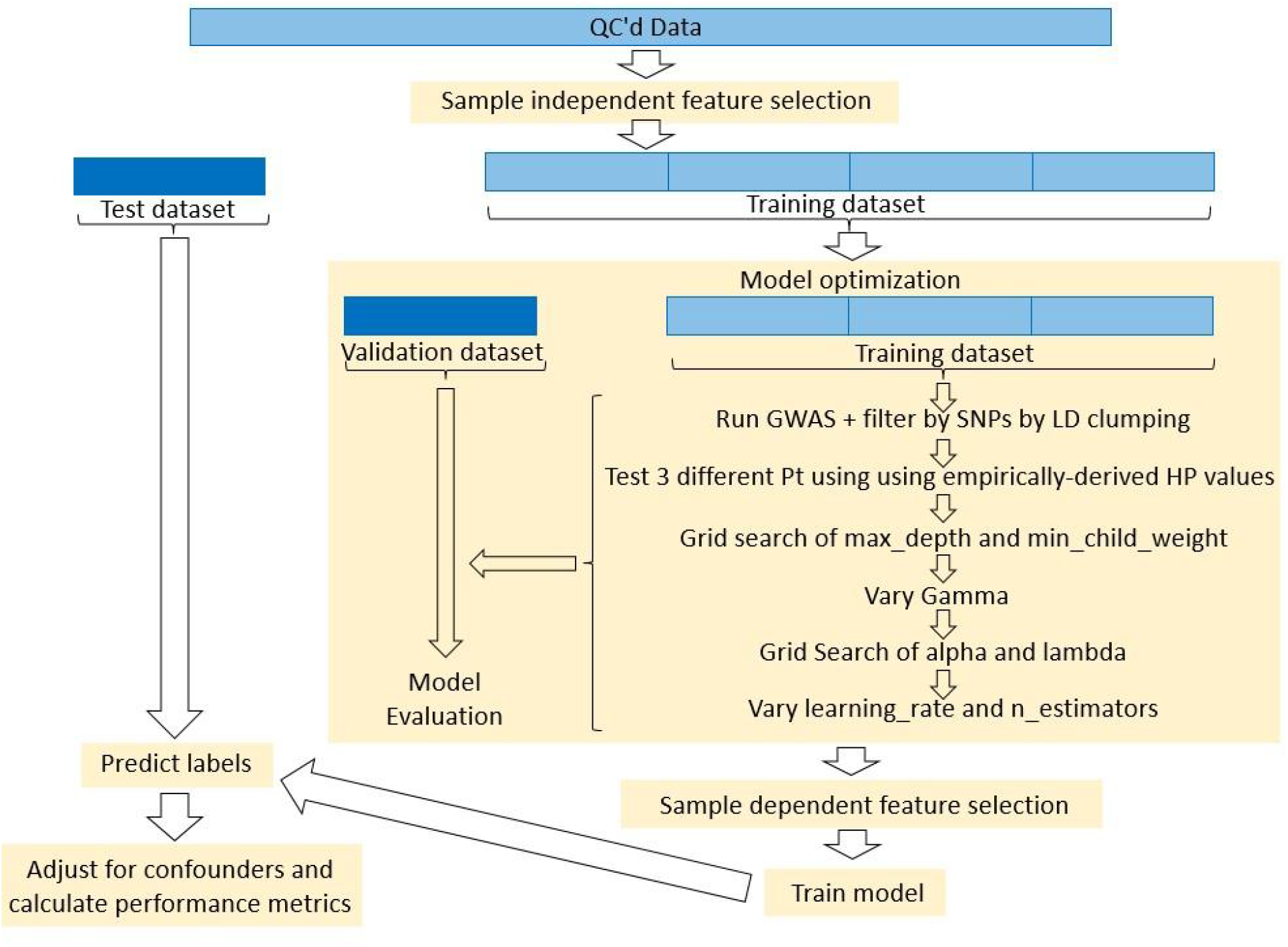
Visualization of the workflow used in this study. QC’d data was first filtered by SNP annotation which is a sample independent step. Next, the data was split into five folds. One of these folds was held out to report the final performance of the model and the remaining four folds were used for model optimisation and training. Model optimization was performed using three-fold cross-validation. In each cross-validation round, one fold was used as validation data to evaluate the models performance and the remaining three folds were used for model training. The first step of model optimization was to run a GWAS on each fold and use the results to filter SNPs by LD clumping. The first model parameter optimized was the p-value threshold (Pt) for feature selection, followed by the XGBoost hyperparameters as outlined in the figure. Model evaluation involved choosing the model with the hyperparameter values that resulted in the highest area under the ROC curve (AUC) value on the validation dataset. After model optimisation, sample dependent feature selection (FS) was performed (this included running a GWAS, performing LD clumping and retaining SNPs that pass the best performing Pt) and the model is run using the best performing hyperparameters on all four training folds. Predictions are made on the held-out test fold. Finally, confounder adjustment is performed post-hoc on the predictions and performance metrics are calculated.

Next, we tuned the first of the XGBoost hyperparameters, min_child_weight and max_depth. These two hyperparameters were tuned together because they both control tree depth. We initially implemented a grid search of three values each for min_child_weight and max_depth, where an XGB model ws run for every combination of hyperparameter values (3 × 3 = 9 different combinations) (Feurer and Hutter, 2019). Based on the results from this grid search, we tested further values for each hyperparameter (results presented in Supplementary Table 1). Next we tuned gamma, followed by the regularization parameters, alpha and lambda. Whereas, the number of trees for p-value optimisation was set at the relatively high value of 15,000, we restricted the number of trees to 6000 for optimization of max_depth, min_child_weight, gamma, alpha and lamda. The reason for this choice was that in initial experiments, we found it possible to identify values for these hyperparameters that provide good performance early on in model building, allowing us to build fewer trees, reducing compute time and memory usage.

Finally, we varied the learning rate and early_stopping_rounds together, as they both control the number of trees in the model. We set the number of trees very high at 35000, to give the model the opportunity to build as many trees as necessary, whilst implementing early_stopping (the model stops building trees if, after a certain number of rounds, no improvement in performance is observed). This approach also helps to prevent overfitting of the model.

The colsampleby family of hyperparameters (colsample_bytree, colsample_bylevel, colsample_bynode) were each set to 0.5. This fraction was chosen to prevent overfitting without being too low as to potentially hinder performance. All other hyperparameters were left at their default values. After optimization, the model was run on the full training dataset using the tuned parameters. Predictions were made on the held-out test set and performance metrics calculated using scikit-learn.

XGBoost was implemented on a HPC system with 2× 20-core 2.4 GHz CPUs and 192 GiB of RAM. There was a time limit (max walltime) of 144 hours for each job. The final job (i.e. training on full dataset and testing on the held out) took 19 hours and 22 minutes to run, using all 40 CPUs. Each run of XGBoost for hyperparameter optimization varied depending on the hyperparameter combinations, particularly the number of trees (between 2 and 17 hours using 40 CPUs). XGBoost was run a total of 96 times for hyperparameter optimisation and once for the final training and prediction of the held out test fold (Figure 2). Code used is freely available at https://github.com/laurafahey02/XGBoost4IntelligencePrediction.

### 2.5 Confounder Adjustment for XGBoost

Output predictions were controlled for the effects of confounders using the method proposed by Dinga *et al*., 2020. This approach involves performing three logistic regressions, where the true test labels are regressed on (1) the confounders and machine learning predictions (full model), (2) just the confounders, and (3) just the machine learning predictions. The amount of variance explained by only the confounds (R^2^_confounders_), variance explained by only the XGBoost predictions (R^2^_xgb_predictions_) and variance not explained by either the confounds or XGBoost predictions (considered the variance explained by both the confounders and the XGBoost predictions; R^2^_shared_) are calculated using simple subtraction equations as described in Dinga and colleagues paper. Nagelkerke’s R^2^ was used to measure variance explained as this is also used by the tool we chose for PGS analysis (detailed in section 3.6), and thus enabled accurate comparison between the two approaches. To take into account non-linear effects of confounds, the splines R package, version 3.6.2, was used to perform cubic spine expansion with 5 knots.

We adjusted for the confounds, age when attended assessment centre (UKB data-field ID 21003), sex, top 8 European principal components, genotyping array, socioeconomic status (assessed by townsend index) and assessment centre that the test was performed in.

### 2.6 Polygenic score (PGS) analysis

The same training and test subsets used for XGBoost were used for PGS analysis. A GWAS was performed on the training subset using logistic regression with firth fallback in PLINK2 and accounting for the same confounders as for the XGBoost model. The resulting summary statistics were used as input for the Polygenic Risk Score software, PRSice-2 (Choi and O'Reilly, 2019), which calculates a PGS for each individual in the test subset. PRSice-2 then regresses each PGS on the true test label and outputs performance metrics (Nagelkerke’s R^2^ and p-value). The same confounders accounted for in the GWAS and XGBoost were adjusted for in this regression.

There is only one hyperparameter required to be optimized for PRSice, and that is the p-value threshold. By default, PRSice tests a large number of p-value thresholds to identify the value that performs the best. It starts at P = 5 × 10^-8^ and increases in intervals of 5 × 10^-5^ up to P = 1. We implemented optimisation using these default settings in a three-fold CV loop. We generated receiver operating characteristic (ROC) curves and calculated the area under the ROC curves (AUCs) using scikit-learn in Python.

### 2.7 Gene Ontology (GO) Over-representation Analysis

SNPs were mapped to HGNC symbols using the ensembl Variant Effect Predictor for GRCh37 using default settings (http://grch37.ensembl.org/Homo_sapiens/Tools/VEP; (McLaren *et al*., 2016). GO overrepresentation analysis was performed using the meta Database ConsensusPathDB web interface (http://cpdb.molgen.mpg.de/). This tool calculates p-values by using a hypergeometric test using all HGNC symbols that are present in at least one GO category as background. All GO categories (biological process, molecular function and cellular component) and levels (1-5) were investigated. Q values were calculated using the false discovery rate method (Benjamini and Hochberg, 1995). This tool was chosen due to its ease of use and the fact that its databases are regularly updated.

## 3 Results

### 3.1 Model Performance

The AUC for the XGBoost model was slightly higher than the AUC for the PGS model (0.644 vs 0.639; Figure 3). However, this may be due to confounders being included in the GWAS logistic regression and not in the XGBoost model. The XGBoost full model (XGBoost predictions and confounders) regressed on the true phenotype labels of the test set resulted in an R^2^ value of 0.139 (p = 3.8 × 10^-253^). This was partly explained by confounders (R^2^_confounsders_ = 0.057 and R^2^_shared_ = 0.022). However, the XGBoost predictions explained a proportion of variance that could not be explained by the effects of confounds (R^2^_xgb_predictions_ = 0.057; Figure 4).

**Figure 3:**
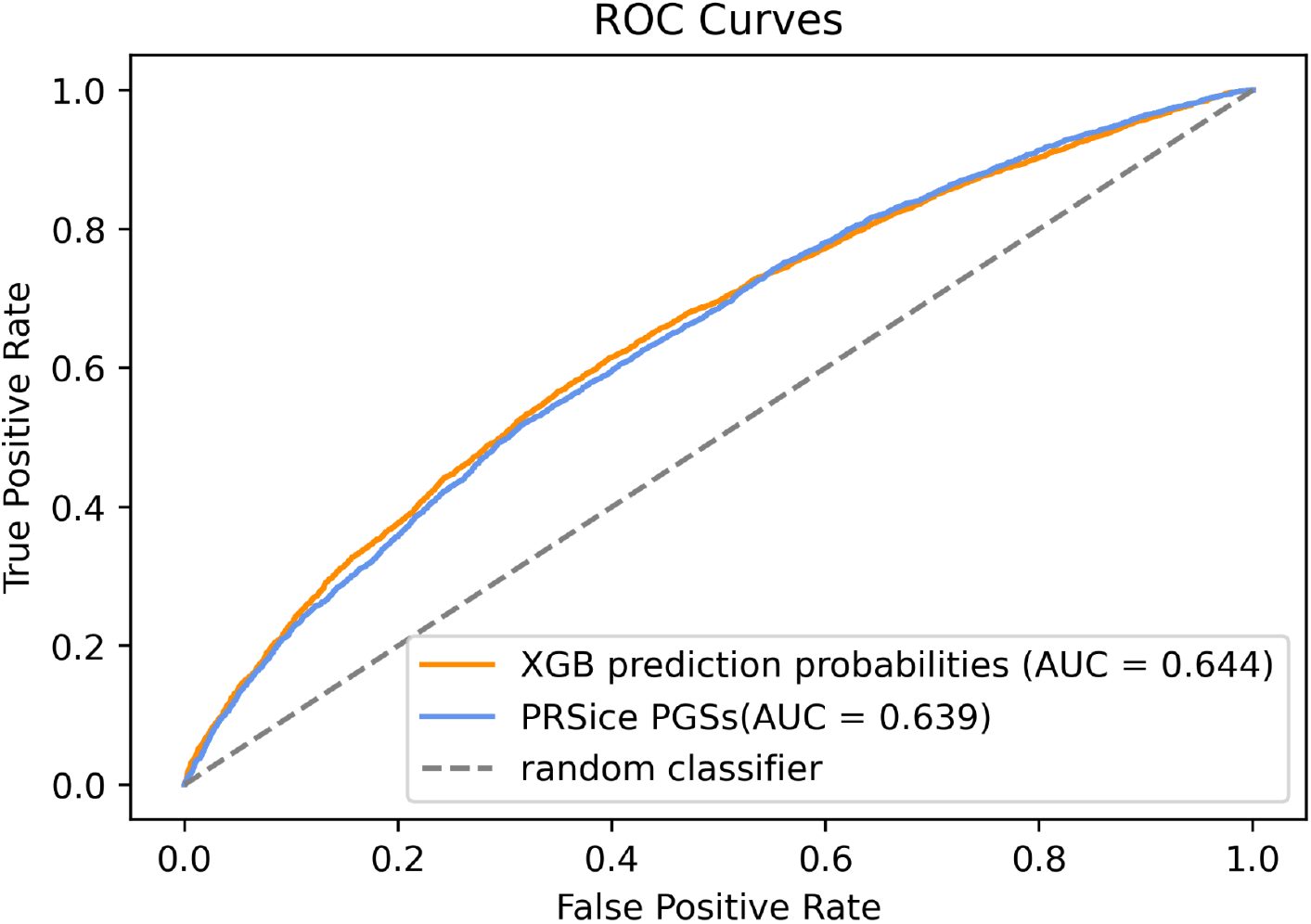
ROC curves for XGBoost (XGB) prediction probabilities and PRSice polygenic score (PGSs).

**Figure 4:**
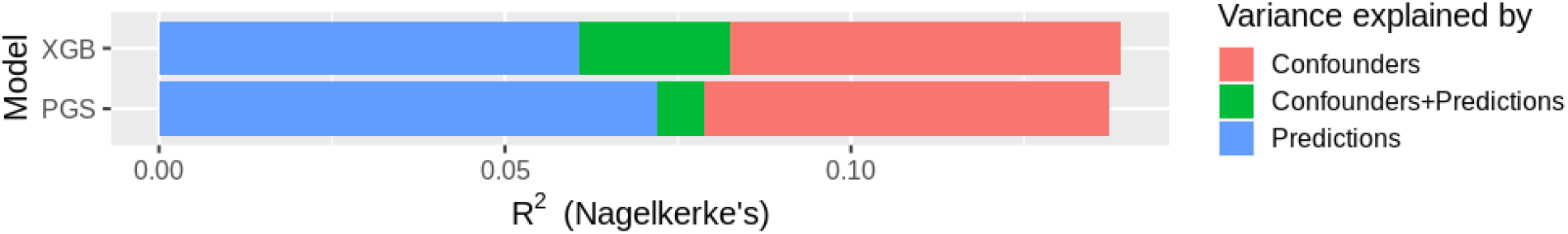
Results from XGBoost (XGB) and polygenic score (PGS) analysis. The XGB full model was able to explain a higher proportion of variance in verbal and numerical reasoning test scores than the PGS full model. However, the variance explained by only the predictions (blue) is lower for XGB than for PGS. This is largely due to a higher amount of shared variance (green) for XGB than PGS.

In comparison, the PRSice full model (polygenic risk scores + confounders) regressed on the true phenotype labels of the test set resulted in the slightly lower R^2^ of 0.137 (p = 2.1×10^-250^). Again, this was partly explained by confounders (R^2^_confounders_ = 0.059 and R^2^_shared_ = 0.007), but to a lesser extent, meaning that the PGS predictions explained a greater proportion of variance that could not be explained by the effects of confounds (R^2^_pgs_predictions_ = 0.072) than the XGBoost predictions (Figure 4).

### 3.2 Best Hyperparameters and Structure of the XGBoost Tree

For PRSice, a p-value threshold of 0.759 was found to perform best. This resulted in 142,584 SNPs being included in PGS construction. For XGBoost, a p-value threshold of 0.2 (second highest value tested) performed best (Table 2). This resulted in 57,381 SNPs input into the model. XGBoost internal feature selection brought this number down to 32,830 SNPs that were used by the model.

The best values for max_depth and min_child_weight were found to be 12 and 150, respectively, meaning performance was best when some level of interaction was modeled between features. The optimal value for gamma was 0.2, meaning a low amount of tree pruning was best. For alpha and lambda, the defaults of 0 and 1, respectively, performed best, indicating a low amount of regularization was optimal. A learning rate of 0.01 performed best and the optimal number of trees was 23,493 (the average number of trees the model grew [with early stopping] across the 3 CV folds; Supplementary Table 1). This high value for the number of trees was expected in order for XGBoost to model the large number of low effect predictors.

To investigate the structure of the XGBoost model, we obtained the count of internal node splits for all trees in the model (N = 423,755) and visualized a tree (Figure 5). Since each node represents a feature which splits the data, and only 32,830 SNPs were used by XGBoost, SNPs were used multiple times by the model. Each tree had an average of 18 node splits per tree (423755 internal splits / 23493 trees).

**Figure 5:**
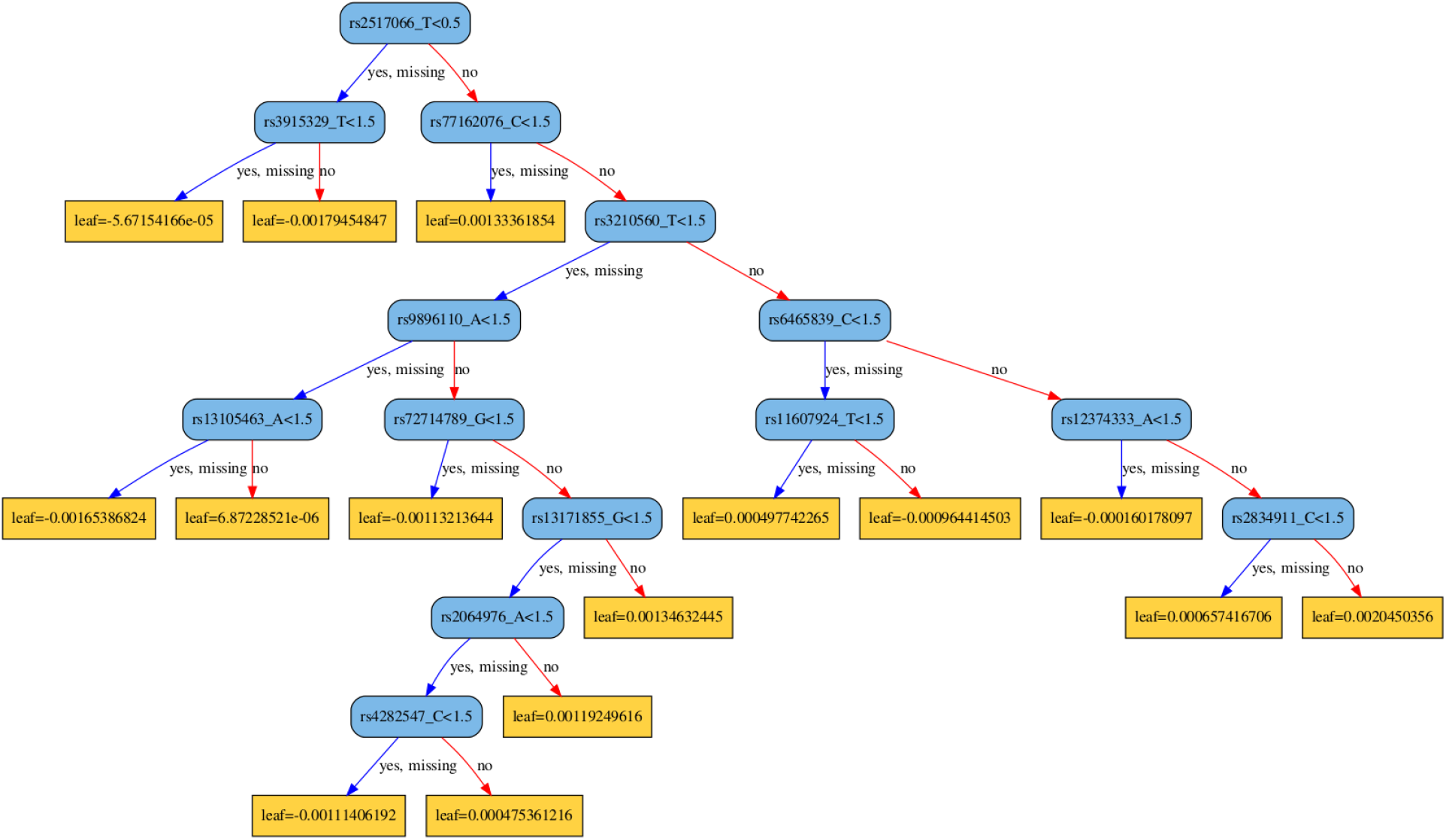
Structure of a tree with 18 splits from our XGBoost model. Nodes are coloured blue and leaves coloured yellow. Each node contains the rsID of a SNP to split the data on. For example, in the root node, the binary question is: is the genotype at rs2517066 less than 0.5. Samples that have a genotype of 0 (no minor alleles) will go to the branch on the left and samples that have a genotype of 1 or 2 will go to the branch on the right. The leaf nodes contain a score for all individuals in that leaf. After all trees have been built, the scores in the leaves of each tree are summed up for each individual and converted to a probability of belonging to the class “HIGH” or “LOW”.

### 3.3 Top Features for XGBoost vs GWAS+PRSice

The most common metric for measuring feature importance in XGBoost is the average gain across all trees. Gain measures the increase in accuracy brought to the branches of a tree by splitting based on that feature. The higher the value for gain, the more important that feature is for prediction. To investigate the relationship between SNPs deemed important by XGBoost, measured using the average gain metric, and SNPs deemed important by GWAS logistic regression, measured using the p-value, we calculated the Pearson correlation between these two metrics for all SNPs used by the XGBoost model. We observed a very low negative correlation with a Pearson correlation of −0.116 (a negative correlation was expected since the higher the gain value the better, whereas the lower the p-value the better). This negative correlation gets stronger when restricting to SNPs with a lower p-value from the GWAS. For SNPs with a p-value < 0.01 (N SNPs = 5,275) the Pearson correlation was −0.209 and for SNPs with a p-value < 0.001 (N SNPs =311) the Pearson correlation was −0.233.

We next investigated differences in gene ontology (GO) enrichment between SNPs deemed important by XGBoost and GWAS. We restricted SNPs to those nominally significant in the GWAS (p-value threshold = 0.05, N SNPs = 14,906) and then extracted the same number of SNPs sorted by gain value to obtain SNPs deemed important by XGBoost. The 14,906 SNPs deemed important by GWAS and XGBoost mapped to 9,929 and 9,796 genes, respectively. In comparison to the low correlation between SNPs deemed important by both models, 70% of these genes overlap. As expected, given this high overlap, the over-represented GO terms are very similar for both sets of genes (Figure 6). The top biological process for both is nervous system development and the top cellular component term for both is neuron part.

**Figure 6:**
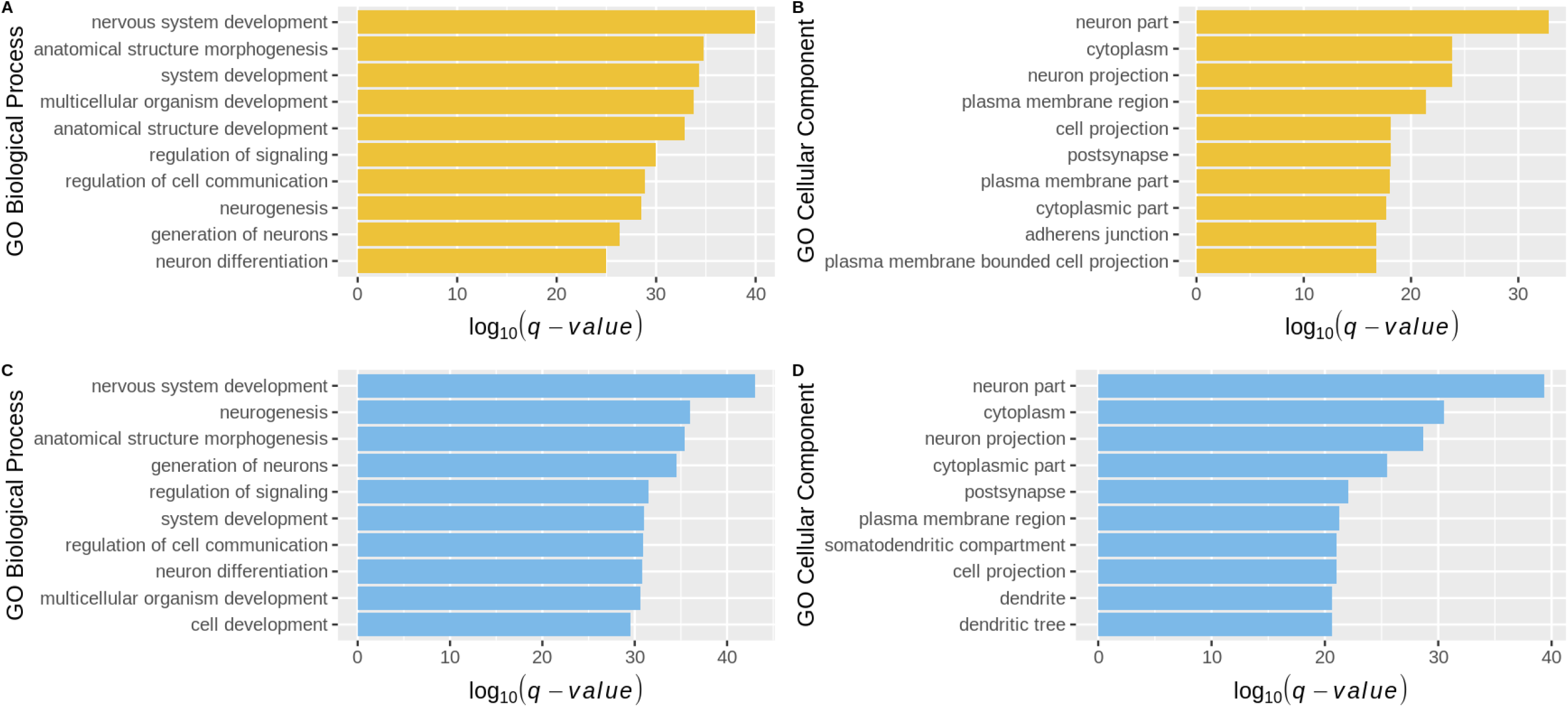
Top 10 GO terms for genes deemed important by both models. For the top 14,906 genes deemed important by XGBoost (A-B; yellow) and PGS (C-D; blue), bar charts visualize the top 10 enriched biological process (A & C) and cellular component (B & D) GO terms

## 4 Discussion

The AUC was higher for XGBoost than for PGS. However, when taking into account the effect of confounding variables, the PGS predictions explained more variation than the XGBoost predictions. The fact that XGBoost performed almost as well as the PGS approach confirms that XGBoost is capable of detecting small effects in complex, polygenic phenotypes. Both GWAS and XGBoost identified similar genes as being important for the prediction of intelligence; 70% of the top ∼9k genes from both methods overlapped, and they both showed enrichment in similar gene ontology categories. A higher p-value threshold was optimal for PRSice (p < 0.76) in comparison to XGBoost (p < 0.2). This resulted in over twice the number of SNPs being used as input into the PGS model in comparison to the XGBoost model (142,584 vs. 57,381 SNPs, respectively). Furthermore, XGBoost performs internal feature selection on the input SNPs which brought the number of SNPs down to 32,830. These 32,830 SNPs were capable of explaining almost the same amount of variation as the 142,584 SNPs required as input for the PGS model. A major challenge of interpreting GWAS results from complex polygenic phenotypes is that there are so many significantly associated variants genome-wide and this makes identifying contributing biological pathways difficult. Although we focused on the top SNPs for our biological interpretation, the fact that the XGBoost model is able to explain the same amount of variation in phenotype as the PGS model with less than a quarter of the SNPs has the potential to aid in biological interpretation of the contributing variants for future studies.

The top SNP identified by XGBoost was rs8084625. This maps to the *Gprotein subunit alpha L* (*GNAL*) gene. GNAL is highly expressed in the central nervous system (CNS). It encodes a subunit of the G protein receptor which mediates odorant signaling. It is located in a susceptibility region for bipolar disorder and schizophrenia, and has been linked to altered epigenetic regulation in SCZ (Corradi *et al*., 2005). This is interesting because cognitive deficits are observed in both of these disorders. A SNP that maps to this gene in the GWAS we performed for PGS analysis, rs1786573, has a p-value of 7.37016×10^-06^, bordering on genome-wide significance. The next top SNP identified by XGBoost, rs118137629, maps to the gene, *extracellular leucine rich repeat and fibronectin type III domain containing 1* (*ELFNI*). ELFNI is also highly expressed in the CNS and encodes a postsynaptic protein. This gene has been found to be associated with mathematical ability (Lee *et al*., 2018), as well as noncognitive aspects of educational attainment in a GWAS by subtraction (Demange *et al*., 2021). SNPs that map to this gene in our GWAS have p-values of 0.004 (rs4720942) and 0.01 (rs10950352). The third top SNP, rs2519757, maps to *TSC Complex Subunit 1* (*TSC1*), which encodes a protein that acts as a tumor suppressor (prevents cells from dividing too much) through negative regulation of the mechanistic target of rapamycin (mTOR) signaling pathway. This pathway has a critical role in neurodevelopment (Cotter, 2020). SNPs that mapped to mTOR in our GWAS had p-values greater than 0.05. The forth top SNP, rs7587835, maps to the gene, *Delta/Notch Like EGF Repeat Containing* (*DNER*). This gene encodes a transmembrane protein and is strongly expressed in Purkinje cells in the cerebellum. DNER functions as a neuron-specific Notch ligand and mediates signaling during glia development through neuron-glia interactions (Eiraku *et al*., 2005). The SNP, rs7587835 from our GWAS mapped to this gene and had a p-value of 0.02. Finally, the fifth SNP, rs5030784, maps to *D-Amino acid oxidase* (*DAO*). Increased expression and activity of the enzyme product of this gene have been reported in the postmortem cerebellar tissue of patients with SCZ (Verrall *et al*., 2007). SNPs in this gene have been reported to be associated with general cognitive ability (p = 7 × 10^-6^) and exploratory eye movement dysfunction in schizophrenia (p = 2 × 10^-7^) through GWAS. In conclusion, the top five SNPs identified by XGBoost are biologically relevant to cognition and not all of these genes were identified as important by the GWAS on the same data, highlighting the fact that examining SNPs in isolation will potentially miss their biological contribution to a complex phenotype.

Scoring interactions as well as individual SNPs may be helpful in order to understand the reason for certain SNPs ranking higher in XGBoost than in the GWAS. For example, a SNP may achieve a higher score in XGBoost than in the GWAS because its effect is dependent on an interaction with another SNP, which would not be captured in the GWAS. A recent study (*Johnsen *et al*., 2021*) used SHapley Additive exPlanation (SHAP) values to score the pairwise contribution of SNPs to XGBoost predictions (rather than using gain values to score the individual contribution of each SNP). However, more work needs to be done to assess the validity of SHap-values in this context (*Johnsen *et al*., 2021*).

Of the previous studies referenced in our introduction that have compared the performance of different ML methods to PGS analysis, only Romagnoni and colleagues achieved the same result as us, that PGS compares similarly to the ML methods tested (Romagnoni *et al*., 2019). Two of the studies found that the ML model outperformed the PGS approach. The first of these, (Behravan *et al*., 2018) used XGBoost, not as a classifier, but as a feature selection method to identify important interacting SNPs (through an adaptive iterative SNP search) to be used as input for a support vector machine classifier. This extra effort to identify interactions possibly contributed to the increased performance of the ML method in comparison to the PGS method. However, this study did not account for confounders in the ML model and also suffered from low sample size. Pinto and colleagues also found the ML method, an artificial neural network, to outperform the PGS in predicting schizophrenia cases from controls, but only for very large sample sizes (N > 30,000) (Pinto *et al*., 2019). This study also did not account for confounders. The Coronary Artery Disease study (Gola *et al*., 2020) was the only study to find the PGS to outperform all ML methods tested. However, the suboptimal performance of the ML methods may have been due to the hyperparameter optimization workflow used. To overcome the computational challenge of hyperparameter optimization, they reduced the outer training fold to a random 20% of samples. Given that the number of samples was not high to begin with (15,709 in the full dataset), 20% of samples may have been too low for adequate hyperparameter tuning.

However, there are also limitations to our current study. Hyperparameter optimization is highly time intensive due to having to run multiple models for different combinations of hyperparameter values; this is further augmented by the requirement of cross-validation to ensure the optimized hyperparameters generalize across the full dataset. The stepwise hyperparameter optimization method we employed here was chosen as it enabled us to search a reasonably large area of the parameter space while maintaining a relatively manageable computational cost. Systematic methods that search through an even greater range of hyperparameter values would potentially discover more optimal values (albeit at increased computational cost).

The number of folds for cross validation in hyperparameter optimization should be large enough so that the best hyperparameters generalize well for multiple data subsets, but not so large that the validation subset ends up being too small to be representative. We chose three CV folds, which should be sufficient to ensure that the best hyperparameters would generalize across the full dataset, but not so large that it would be too computationally costly. It is, however, possible that a higher number of folds would have led to better hyperparameter values. Outer cross validation of the training and held out test set was not performed as it was too computationally intensive to run the whole workflow multiple times. However, we believe the sample size used here was large enough (46,047 training samples and 11,513 test samples) that this was not an issue.

Feature selection in this study involved filtering by SNP annotation, LD clumping and p-value thresholding, prior to being input into the XGBoost model. There exists multiple filter methods that can capture nonlinear interactions between features; popular methods include Relief-based algorithms (Urbanowicz *et al*., 2018) and minimum redundancy maximum relevance (mRMR; Ding and Peng, 2005). It is possible that capturing nonlinear interactions in the filtering step would have led to a greater final performance. However, current methods are too time and memory intensive to be feasibly implemented on very large genomic datasets.

In conclusion, we found that XGBoost is capable of detecting small SNP effects for the complex polygenic phenotype, intelligence, assessed using the UKB verbal and numerical reasoning test. Although XGBoost did not outperform the classical PGS method, it was capable of achieving good predictive performance with a fraction of the SNPs and the top SNPs identified as being important for predictive performance were biologically relevant.

