## Supplementary Material for "Comparing the XGBoost machine learning algorithm to polygenic scoring for the prediction of intelligence based on genotype data"

### UK Biobank Data Quality control

We restricted samples to those of European ancestry. Such samples were identified using principal component analysis (PCA) and 1000 Genomes Project (1KGP) data. This was done by first merging 1KGP variant call format (VCF) files with UK Biobank (UKB) directly genotyped PLINK files. SNPs were restricted to only those used for PCA by UKB (identified using the UKB supplied marker QC file). PCA was performed on this merged set of PLINK files in PLINK2 (www.cog-genomics.org/plink/2.0/; Chang et al., 2015) using the --approx option to reduce memory requirements. The multi-mean of the top eight PCs for 1KGP samples of European ancestry (identified via CEU code) was calculated and UKB samples with a Mahalanobis distance < 6 SD from this multi-mean were identified as being of European ancestry.

Further samples were removed on the basis of high SNP missingness, unusually low of high heterzygosity, discordent sex informaiton, and containing chromosomal aneuploidies. These samples were identified by UKB (Bycroft et al. 2018) and listed in the UKB supplied sample QC file. We next removed related samples using the UKB supplied relatedness file which lists pairs of individuals related up to the third degree. One individual was removed from each pair, preferentially keeping samples with Fluid Intelligence phenotype data and skipping pairs with one member already removed. The total number of samples after QC was 333,706. The phenotype used for this study was UK Biobank (UKB) Fluid intelligence score (Data-field ID 20016). We used UKB Fluid intelligence scores collected at the initial assessment visit. The number of samples with phenotype data post QC was 118,021.

Imputed variants were converted to hard calls at a certainty threshold of 0.9. SNPs were excluded if their proportion of missing genotypes exceeded 2%, minor allele frequency (MAF) was less than 1%, or Hardy–Weinberg equilibrium (HWE) was lower than $1\times10^{-6}$. These three metrics were calculated in PLINK2. Multi-allelic snps and SNPs with differences in allele frequency between the two genotyping arrays, used were also removed. The total number of SNPs after QC was 7,832,917.

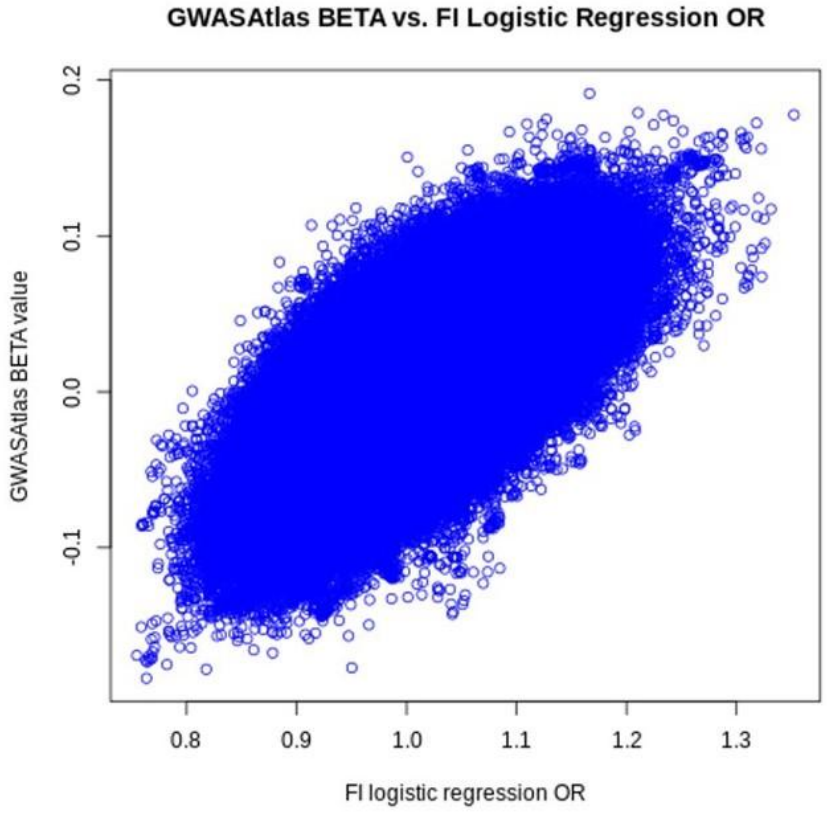

**Supplementary Figure 1: GWASAtlas linear regression BETA value vs. logistic regression odds ratio (OR)**. Correlation between the beta values from the linear regression GWAS performed on the full quantitative fluid intelligence (FI) phenotype by GWASAtlas and the odds ratios (OR) from the logistic regression GWAS we performed on the binary FI phenotype. Only SNPs common to both summary statistic files were plotted.

**Supplementary Table 1: Average area under the curve (AUC) and number of trees across the three cross-validation folds of all combinations of hyperparameters and their values tested.**

| Hyperparameter(s) | Value(s) | Average AUC | No. trees |
|---|---|---|---|
| p-value threshold | 5x10-5 | 0.5478429 | 15000 |
| | 0.05 | 0.6046115 | 15000 |
| | 0.1 | 0.6073059 | 15000 |
| | 0.2 | 0.6087605 | 15000 |
| | 0.3 | 0.6059531 | 15000 |
| max_depth & min_child_weight | 4 & 10 | 0.5846155 | 6000 |
| | 4 & 50 | 0.5955235 | 6000 |
| | 4 & 100 | 0.5990083 | 6000 |
| | 8 & 10 | 0.5991444 | 6000 |
| | 8 & 50 | 0.6029171 | 6000 |
| | 8 & 100 | 0.6062539 | 6000 |
| | 8 & 150 | 0.6061069 | 6000 |
| | 8 & 200 | 0.604566 | 6000 |
| | 12 & 10 | 0.5981851 | 6000 |
| | 12 & 50 | 0.6016336 | 6000 |
| | 12 & 100 | 0.606491 | 6000 |
| | 12 & 150 | 0.6075969 | 6000 |
| | 12 & 200 | 0.6045559 | 6000 |
| | 14 & 100 | 0.6044329 | 6000 |
| | 14 & 150 | 0.6043446 | 6000 |
| | 14 & 200 | 0.6045559 | 6000 |

*To be continued.*

**Supplementary Table 1 (continued)**

| **Hyperparameter(s)** | **Value(s)** | **Average AUC** | **No. trees** |
|---|---|---|---|
| gamma | 0.1 | 0.6081661 | 6000 |
| | 0.2 | 0.6090466 | 6000 |
| | 0.3 | 0.6088096 | 6000 |
| | 0.4 | 0.6070636 | 6000 |
| | 1 | 0.6044558 | 6000 |
| alpha & lambda | 0 & 0.1 | 0.6067992 | 6000 |
| | 0 & 1 | 0.6090466 | 6000 |
| | 0 & 5 | 0.6053998 | 6000 |
| | 0.01 & 0.1 | 0.6033153 | 6000 |
| | 0.01 & 1 | 0.6045318 | 6000 |
| | 0.01 & 5 | 0.6053505 | 6000 |
| | 0.05 & 0.1 | 0.6043953 | 6000 |
| | 0.05 & 1 | 0.6054501 | 6000 |
| | 0.05 & 5 | 0.6053452 | 6000 |
| num_boost_round & learning_rate | 100 & 0.05 | 0.5948518 | 1874 |
| | 1000 & 0.05 | 0.6126017 | 12304 |
| | 1000 & 0.01 | 0.6173017 | 15817 |
| | 1500 & 0.01 | 0.6186966 | 23493 |
| | 2000 & 0.01 | 0.6173017 | 16816 |
| | 1500 & 0.005 | 0.6164639 | 18365 |

*Note: Best performing HP and their corresponding AUC are highlighted in red.*
